# Argon neuroprotection in a non-human primate model of transient endovascular ischemic stroke

**DOI:** 10.1101/2024.01.24.577050

**Authors:** S Gonzalez Torrecilla, A Delbrel, L Giacomino, D Meunier, J Sein, L Renaud, P Brige, P Garrigue, JF Hak, B Guillet, H Brunel, G Farjot, T Brochier, L Velly

## Abstract

**Background:** Previous studies have demonstrated the efficacy of argon neuroprotection in rodent models of cerebral ischemia. The objective of the present study was to confirm a potential neuroprotective effect of argon in a non-human primate model of endovascular ischemic stroke as an essential step before considering the use of argon as a neuroprotective agent in humans.

**Methods:** Thirteen adult monkeys (*Macaca mulatta*) were allocated to two groups: a control group (n=8) without neuroprotection and an argon group (n=5) in which argon inhalation (90 min) was initiated 30 minutes after onset of ischemia. Animals in both groups underwent brain MRI (pre-ischemic) at least 7 days before the intervention. The monkeys were subjected to focal cerebral ischemia induced by a transient (90 min) middle cerebral artery occlusion (tMCAO). After tMCAO, MRI was performed 1 hour after cerebral reperfusion. The ischemic core volume was defined by the apparent diffusion coefficient (aDC) and edema in fluid attenuated inversion recovery (FLAIR) acquisitions. MRI masks were applied to distinguish between cortical and subcortical abnormalities. In addition, a modified version of the Rankin scale was used to neurologically assess post-tMCAO.

**Results:** Despite variability in the ischemic core and edema volumes in the control group, argon significantly reduced ischemic core volume after ischemia compared to the control group (1.1±1.6 cm^3^ *vs.* 8.5±8.1 cm^3^; *p*=0.03). This effect was limited to cortical structures (0.6±1.1 cm^3^ *vs.* 7.4±7.2 cm^3^; *p*=0.03). No significant differences were observed in the edema volumes. Measures of neurological clinical outcome suggested a better prognosis in argon-treated animals.

**Conclusions:** In the tMCAO macaque model, argon induced effective neuroprotective effects, leading to a reduced ischemic core in cortical areas. These results support the potential use of this therapeutic approach for future clinical studies in stroke patients.

## INTRODUCTION

With 5.4 million deaths each year, the World Health Organization (WHO) considers strokes to be the second leading cause of death worldwide[1]. Strokes are also a major cause of adult disability for those surviving the cerebral insult. Eighty percent of strokes have an ischemic origin due to the occlusion of large intracranial arteries[2], impacting the supply of oxygen and other nutrients to the brain[3].

Discovery and development of safe therapeutics agents with neuroprotective potential for ischemic stroke remains a large and promising field of research[2]. Damaged cells in the penumbra of the ischemic core have been one of the priority targets of these approaches, but, until very recently and a Toll-like receptor 4 antagonist[4], most neuroprotective strategies have failed in clinical practice[3]. In the past decade, noble gases have emerged as highly promising neuroprotective agents. Numerous *in vitro* and *in vivo* studies in rodents have demonstrated the neuroprotective potential of these gases in various models of hypoxia and/or ischemia[5,6]. Xenon was initially identified as the most promising agent, but its rarity and excessive production cost preclude its widespread clinical application[7]. However, several *in vitro* have shown argon also offers neuroprotective effects against hypoxic-ischemic events[8]. Although called “inert” gaz, argon may exert certain biological effects, especially interacting with larger proteins, protein cavities or even receptors [9]. There are many pieces missing to complete the signaling pathway throughout the cell, but the proposed mechanisms for argon mediated neuroprotective effects includes Toll-like receptor 2 and 4 signaling[10]. Importantly, unlike xenon, argon inhalation has neither narcotic nor anesthetic actions under normobaric pressure, it doesn’t present the potential of drug-interaction with rTPA, and can be produced in large amount[11,12]. Short-term beneficial effects of argon have also been reported *in vivo* in rats and pigs in various experimental models of cardiac arrest, neonatal hypoxic-ischemic encephalopathy, and retinal ischemia[13]. Moreover, the positive effect of argon neuroprotection on infarct volume has been confirmed in murine models of transient middle cerebral artery occlusion (tMCAO)[12,14].

Although these observations support a neuroprotective effect of argon inhalation, the translational potential of this therapeutic approach for clinical applications remains speculative due to the significant differences between the brains of humans and rodents. In the present study, we evaluated the neuroprotective effect of argon in a non-human primate model (NHP) of tMCAO. Based on MRI investigations, we analyzed the effect of argon inhalation on the ischemic core and edema volumes measured soon after post-ischemic cerebral reperfusion. In addition, we evaluated longitudinal functional outcome.

## METHODS

All experiments were carried out following European Directive 2010/63/UE and approved by the ethics committee C2EA 71 (Aix-Marseille University, authorization APAFIS#6179-2016072212029223V4). The study included 16 male rhesus monkeys with an average age of 9.6±3.2 years and average body weight of 12.8±4.1 kg. Two of the monkeys were involved in a pilot phase to refine the endovascular occlusion protocol and adjust the acquisition parameters for MRI and, therefore, were not included in the analysis. Of the remaining 14 monkeys, two were reused from a previous project and, for ethical reasons, were euthanized at the end of post-ischemic MRI. The other 12 animals were randomly allocated using a random-number table to two groups: a control group without neuroprotection and an argon group in which argon inhalation was initiated 30 minutes after onset of ischemia. Animals in both groups underwent the tMCAO protocol and were used for MRI and post-ischemic neurological assessment (Figure 1A). One animal in argon group was excluded prior to tMCAO due to severe arthritis which prevented training in behavioral tests.

**Figure 1.**
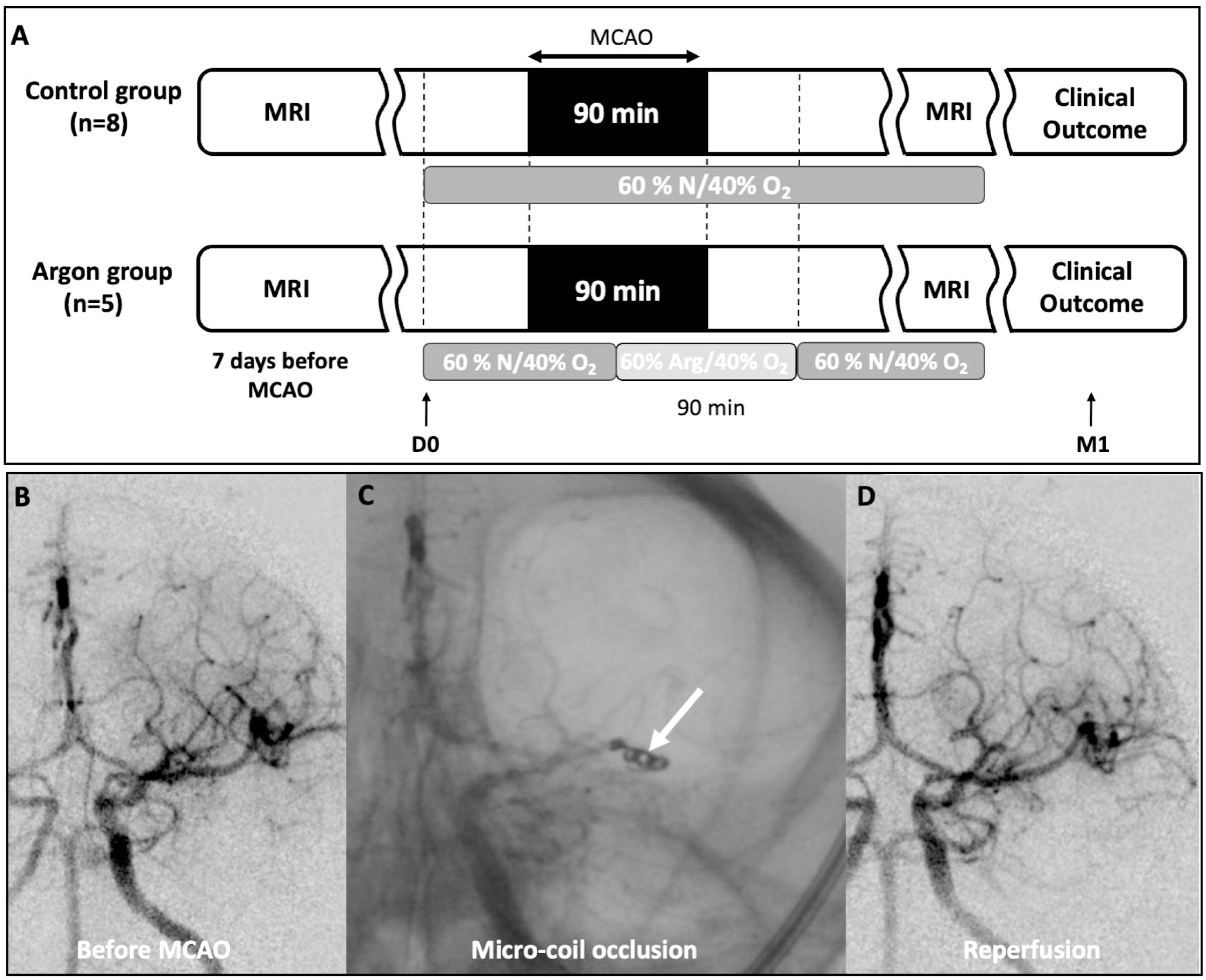
Experimental protocol: (A) CONSORT diagram (B) Project organization. All animals underwent MRI 7 days before the intervention. The animals were then separated into two groups (control and argon) to undergo 90 min tMCAO. For the argon group, argon gas was applied 30 minutes after the onset of ischemia for 90 minutes. (C-E) tMCAO in macaque Rhesus. (C) Cerebral arteriography with contrast agent injected into the right internal carotid before, (D) during, and (E) after MCAO followed by reperfusion. The white arrow indicates the location of a micro-coil occlusion. The total reperfusion of the ischemic territory was carefully controlled after micro-coil removal by angiography. All animals underwent cerebral MRI within one hour of cerebral reperfusion to assess the ischemic core, oedema volumes and confirm the persistence of reperfusion.

The animals were housed in pairs in enriched cages in a temperature- and hygrometer-controlled room with a 12:12 h light-dark cycle. A laboratory diet was provided twice daily, supplemented with fresh fruits and vegetables and *ad libitum* access to water. All procedures were performed under veterinary supervision with the objective to limit pain and discomfort and to promote animal welfare.

### Transient middle cerebral artery occlusion

In all animals, transient cerebral ischemia was achieved by an endovascular approach without craniotomy. Briefly, 24 hours before ischemia, NHPs received anti-platelet aggregation medication (ticagrelor 45 mg) to promote tMCAO recanalization at the end of the procedure. On the day of ischemia, NHPs were sedated by intramuscular injection of ketamine (10 mg/kg), midazolam (0.1 mg/kg), and glycopyrrolate (0.005 mg/kg) and warmed by a heated blanket (Bair Hugger®, 3M, Minnesota, USA) to maintain a body temperature of 38.0±0.5°C throughout the procedure. After an IV line was inserted in the saphenous vein, a bolus injection of propofol (2 mg/kg) was administered to facilitate tracheal intubation (probe diameter 4.5-5 mm). In both groups, after intubation, the lungs were mechanically ventilated (ventilator Zeus, Drager, Lûbeck, Germany) with 2.3 vol% end-tidal sevoflurane (Baxter, Guyancourt, France) in a gas mixture of 40% oxygen and 60% nitrogen (Figure 1B), a tidal volume of 6 mL/kg, and a PEP of 3 cmH_2_O. Ventilation was adjusted to maintain an exhaled CO_2_ level of 30 to 40 mmHg. The animal’s condition was continuously controlled for heart rate, oxygen saturation, capnography, and temperature (Infinity Delta and C700; Dräger, Antony, France). An arterial catheter was placed at the left femoral artery to obtain blood pressure continuously and to facilitate blood sampling during the procedure. A puncture was made in the right femoral artery at the right inguinal fold under ultrasound (Toshiba Aplio XV, Tokyo, Japan), and an experienced interventional neuroradiologist introduced a 4 French catheter (Terumo, Guyancourt, France) to advance it via the aorta into the internal carotid. Angiography (OEC Brivo plus®, GE Healthcare, Buckinghamshire, England) was used to control the real-time catheter positioning (Figure 1C). Heparin 50 units/Kg units IV was given once the microcatheter reaches the middle cerebral artery in desired position. A micro-coil (HyperSoft Helical, 2 mm x 3 cm, TERUMO, Japan) was positioned in the distal M1/proximal M2 portion of the middle cerebral artery to trigger complete occlusion (Figure 1D). Blood gas analyses (PaO_2_, PCO_2_, Hemoglobin, glycemia) were performed during ischemia using the GEM 5000 (Werfen, Le Pré Saint Gervais, France). After a 90-min ischemic interval, the micro-coil was withdrawn, allowing reperfusion. The procedure was considered technically adequate if angiography with an injection of contrast agent confirmed the complete interruption of blood flow at the M2 level (Figure 1D) and the complete reperfusion of the brain after the 90 minutes ischemia (Figure 1E). *In situ* thrombolysis with alteplase (6 mg) was supplied in case of incomplete recanalization and nimodipine in case of vasospasm. At the end of the procedure, hemostasis of the femoral arterial access site was obtained by manual compression.

### Exposure to argon

In the argon group, argon was applied 30 minutes after the onset of ischemia to 30 minutes after reperfusion for a total duration of 90 minutes (Figure 1B). Argon conditioning consisted of exposure to a pre-mixed gas with 40% oxygen and 60% argon (Air Liquide, France). Following argon exposure, the animals were returned to a gas mixture of 40% oxygen and 60% nitrogen and moved for MRI. In contrast, animals in the control group were continuously exposed to a gas mixture of 40% oxygen and 60% nitrogen during the tMCAO procedure and subsequent MRI.

### MRI acquisition

Two MRI scans were performed on each animal. An initial control MRI was performed at least 7 days before the tMCAO. A second MRI was performed on the day of the tMCAO 1 hour after cerebral reperfusion (Figure 1B).

For both MRI sessions, the animal was supine and immobilized with foam in the MRI head coil during scanning. The MRI was performed on a Siemens 3T Prisma MRI scanner (Siemens Medical Solutions, Saint-Denis, France) with a 16-channel receive-only antenna (Scanmed, Omaha, NE, USA) and consisted of a high-resolution T1-weighted structural image with a Magnetization Prepared -RApid Gradient Echo (MPRAGE) three-dimensional protocol (3DT1; slice thickness=0.4 mm; TR/TE= 3300/3.13 ms, FOV=128 mm×128 mm, flip angle=8 degrees, TI=1120 ms, matrix=320×320, GRAPPA=2, 3 averages), a T2-weighted structural image with a Sampling Perfection with Application optimized Contrasts using different flip angle Evolution (SPACE) three-dimensional protocol (3DT2; slice thickness=0.4 mm, TR/TE =3200/544 ms, FOV=128 mm×128 mm, matrix=320×320, 2 averages), diffusion-tensor imaging (DTI; slice thickness=1 mm, single-shot EPI sequence with multiband acceleration factor=2 with TR/TE=3430/82 ms, 92 directions with b-values=0, 300, 1000 s/cm^2^, FOV=112×112 mm, data matrix=112×112; repeated twice for each phase encoding direction, giving a total of 223 volumes per encoding direction; the average diffusivity coefficient [aDC]=[lambda(1)+lambda(2)+lambda(3)]/3 where lambda(1), lambda(2), and lambda(3) are the eigenvalues of the diffusion tensor maps uniformly recreated by our pipeline), an axial 2D T2-weighted fluid-attenuated inversion recovery (FLAIR; slice thickness=1.5 mm, TR/TI/TE=8370/2370/104 ms, FOV=192x132 mm^2^, data matrix=192×132, GRAPPA=2, 2 averages), and time-of-flight magnetic resonance angiography (MRA TOF; FOV=140×192 mm^2^, slice thickness=0.6 mm, TR/TE=21/4.05 ms, 5 slabs, 94 slices, 234x320 matrix, zero-filling interpolation, GRAPPA=2, single average).

### DTI and FLAIR-based assessment of the “ischemic core” and “edema”

To facilitate the extraction of reliable morphometric data and provide a common reference space for group-level analyses, the MRI scans were preprocessed using the software *Macapype* (https://macatools.github.io/macapype/index.html), FSL (http://www.fmrib.ox.ac.uk/fsl), and SPM12 (https://www.fil.ion.ucl.ac.uk/spm/software/spm12/) and aligned on the T1 and normalized in the macaque MRI template INIA19 space (http://nitrc.org/projects/inia19/; Supplementary Figure 1). Two masks were used to separately quantify marked voxels from the cortical or subcortical areas (Supplementary Figure 2). The total volume of one hemisphere in the common INIA19 reference space was 57 cm^3^, subdivided as 43 cm^3^ for the cortical volume and 14 cm^3^ for the subcortical volume.

For each animal, the “ischemic core” (hypointensity on MRI aDC maps) and “edema” (hyperintensity on MRI FLAIR maps) volumes were manually delineated by two trained researchers (GTS, DA) blinded to experimental groups using Freeview software (Figure 2) and average volumes compared between groups (Figure 3). If necessary, images were left-right inverted so that all injuries appeared in the right hemisphere. From normalized and binarized volumes, overlap maps were generated for “ischemic core” and “edema” to show the frequency of injuries on a voxel-by-voxel basis (Figure 4).

**Figure 2.**
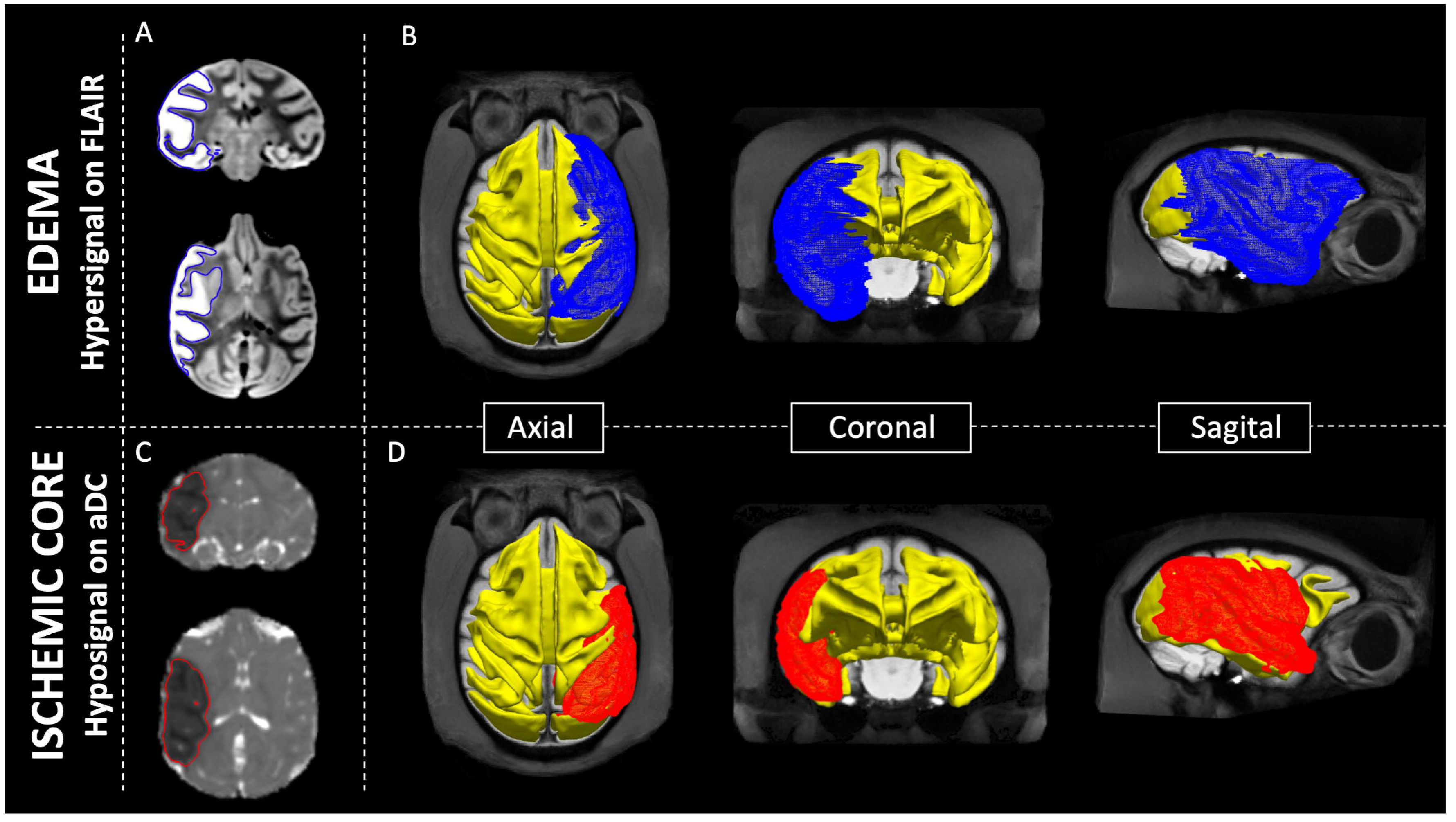
Ischemic core and edema volume reconstructions: (A, C) Pre-processed axial and coronal aDC and FLAIR images for one monkey in the control group following right tMCAO. The ischemic core and edema contours were manually delineated on every axial slice. (A) On the FLAIR, edema appears as hyperintensity. (C) On the aDC, the ischemic core appears as hypointensity. (B, D) Volume reconstruction of the edema (red) and ischemic core (blue) from the aDC and FLAIR images, respectively. All images are from the same subject.

**Figure 3.**
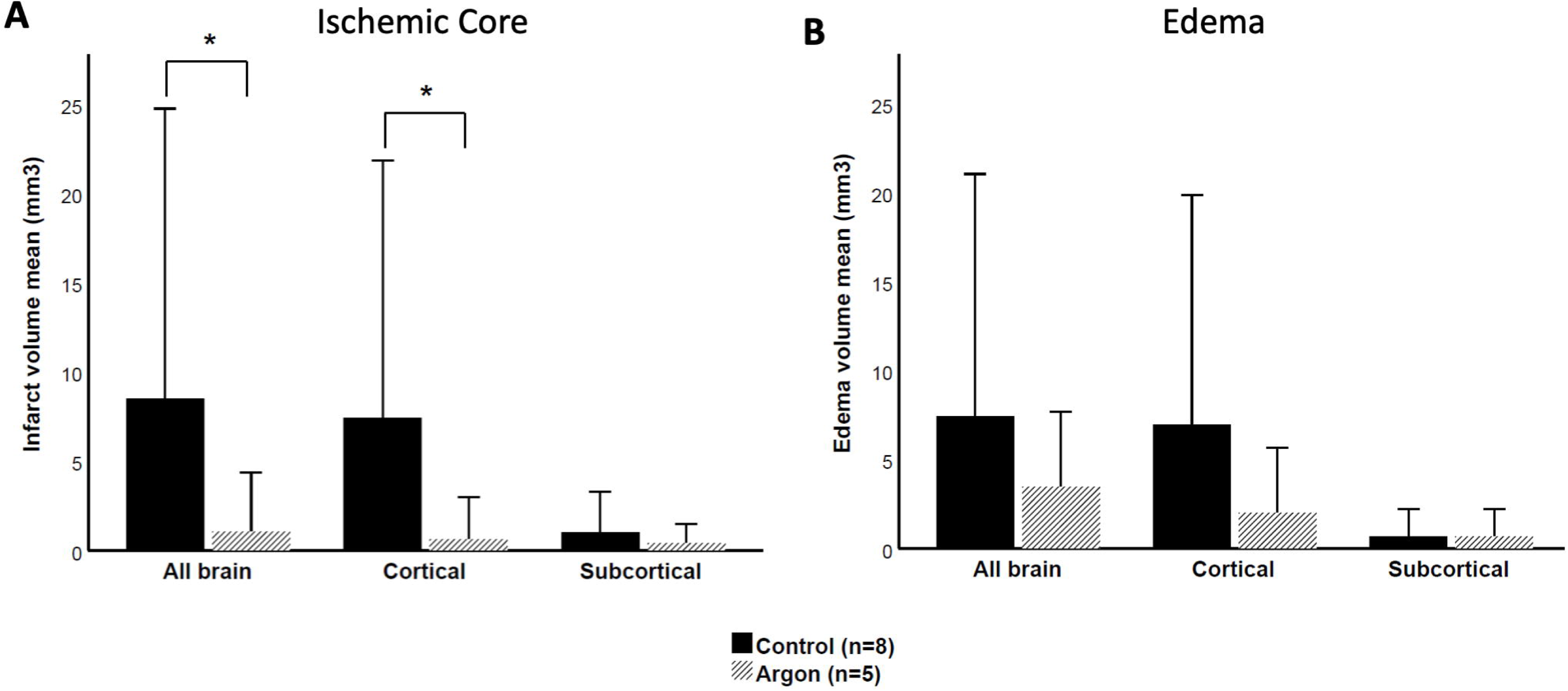
Average ischemic core and edema volumes: (A) The infarct volume was significantly smaller in the argon group than the control group in the entire brain and at the cortical level. The subcortical volumes presented no significant differences between groups. (B) There was no significant difference in the edema volume between groups. Data are presented as mean ± SD. * *p*<0.05 versus control group.

**Figure 4.**
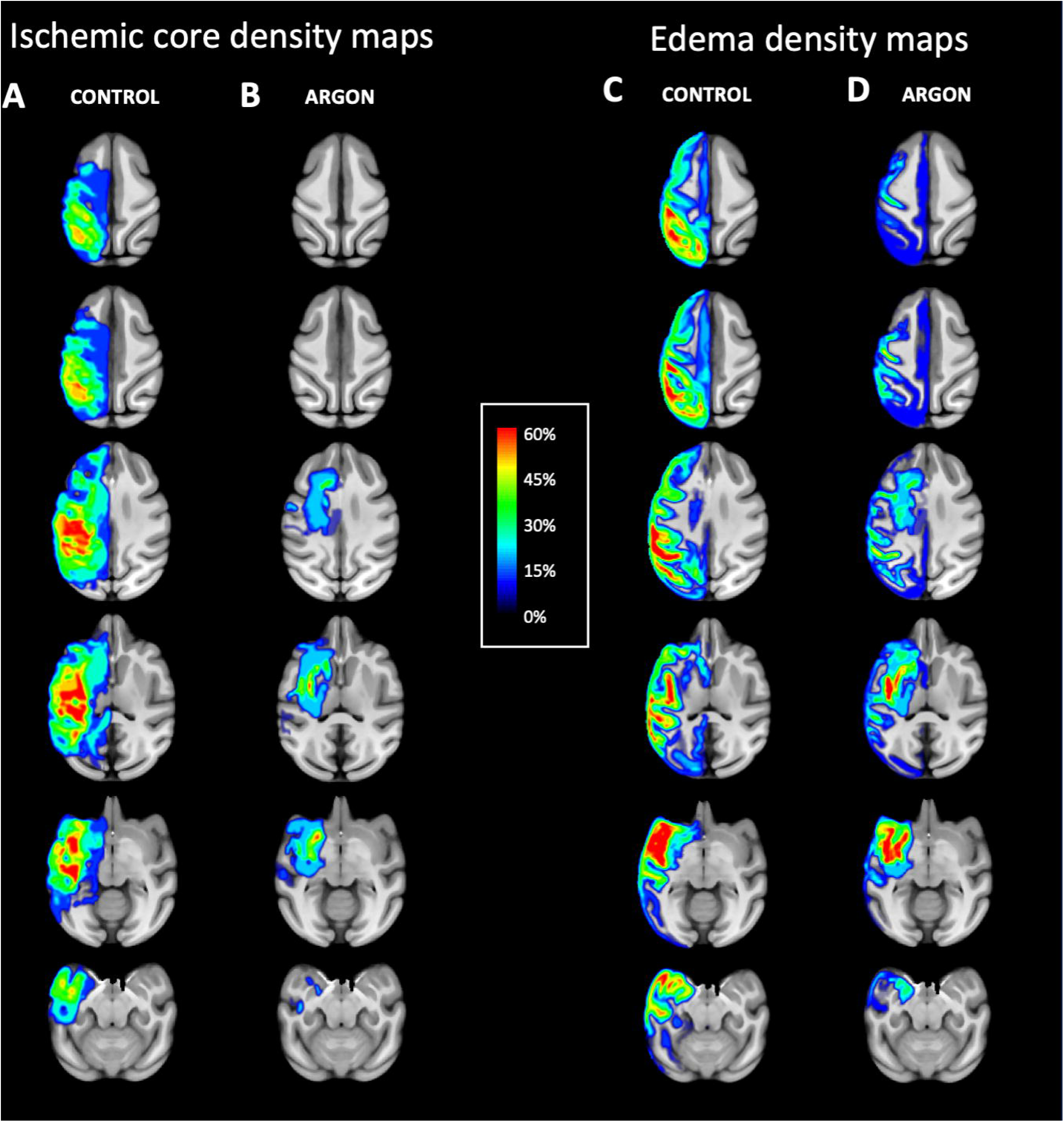
Density maps: (A, B) Overlay plots for the “ischemic core” and (C, D) “edema” computed across all animals of the argon and control groups, respectively. The color overlay created on top of the INIA19 standard brain template indicates the percentage of animals with an ischemic core or edema at each voxel. Warmer colors indicate increasing probability of damage (see color bar).

### Primate Rankin scale

To evaluate the short- and long-term neurological dysfunction in NHP, we used an adapted Rankin scale (mRS). Hours after tMCAO mRS data were collected and for several weeks, we evaluated the monkeys with this scale until they reached a stable neurological state (Figure 5). Based on a previous study[15], we focused our clinical evaluation on hand function and strength, level of activity, and general mobility. A score of 0 indicated no identifiable deficit; 1 indicated reduced use of the upper limb, awkward reach, and reduced strength; 2 an inability to grasp with the affected arm, a slight limp, and reduced self-care; 3 severe disability, in which animals were unable to walk and self-clean without assistance and had limited responses to external stimuli; and 4 indicated the subject’s death.

**Figure 5.**
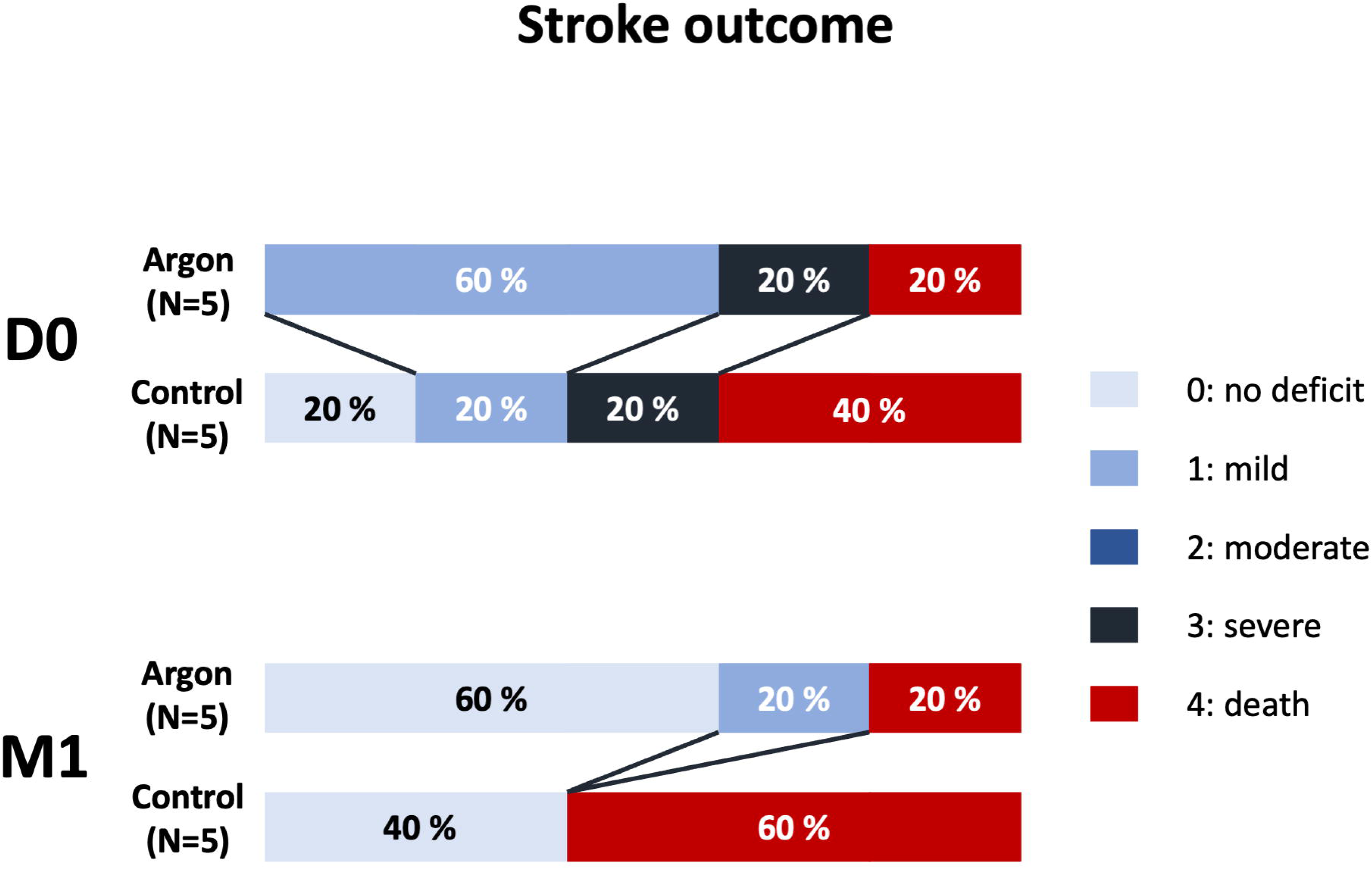
Stroke outcome: Neurological deficits were evaluated on day 0 (DO) and 1 month (M1) after tMCAO using the mRS score (normal score: 0; death: 4). On day 0, the scores were variable in both groups, ranging from no deficit to death (1 and 2 deaths in the argon and control group, respectively). Between-group differences were more prominent at M1, with milder deficits and less death in the argon group compared to the control group.

### Statistical analysis

The number of monkeys per group was assessed by performing a power analysis using the online software (http://www.dssresearch.com/toolkit/sscalc/size_a2.asp). From preliminary in vivo experiments [11] we expected a 40% reduction of cortical infarct volume in the argon group with α error level (p)<0.05; β error level=0.5. This led to a minimum number of animals of n=5 for both samples.

Means and standard deviations were used for normal distributions of continuous variables, and medians and inter-quartile ranges (IQR) for non-continuous variables. For categorical variables, numbers and percentages were used. Differences between continuous variables were assessed by an unpaired two-sample t-test with Bonferroni corrections for *post hoc* intergroup comparisons or a Wilcoxon–Mann–Whitney U test (no assumption for distribution). Comparison between continuous variables from two groups were assessed by Fisher’s exact test. SPSS^®^ Statistics v25.0 software (IBM Corp, Armonk, NY) was used for statistical comparisons. An alpha level of *p*<0.05 was considered significant.

## RESULTS

NHPs were allocated to a control group (n=8) and an argon group (n=5) respectively (Figure 1E). Physiological variables were recorded before, during, and after tMCAO in the 13 animals that participated in the tMCAO protocol. There were no significant differences in age, weight, heart rate, mean arterial pressure, temperature, PCO_2_, PO_2_, hemoglobin or glucose concentration between the control and argon groups (Table 1). *In situ* thrombolysis with alteplase (6 mg) was supplied to four animals (control group n=1; argon group n=3) and nimodipine was supplied to four animals (control group n=3; argon group n=1). The “ischemic core” and “edema” volumes measured in the control (n=8) and argon groups (n=5) are presented in Figure 3. Argon significantly reduced the “ischemic core” compared to the control group (1.1±1.6 vs. 8.5±8.1 cm^3^; *p*=0.03). After applying the binary masks, the difference between the ischemic core of the argon and control groups was still significant at the cortical level (0.6±1.1 vs. 7.4±7.2 cm^3^, *p*=0.03) but not significant at the subcortical surface level (0.4±0.5 vs. 1.0±1.1 cm^3^; *p*=0.09). Non-significant differences were found between the edema volumes in both groups (3.4±2.1 vs. 7.4±6.8 cm^3^; *p*=0.16), even after applying the binary masks.

**Table 1:**
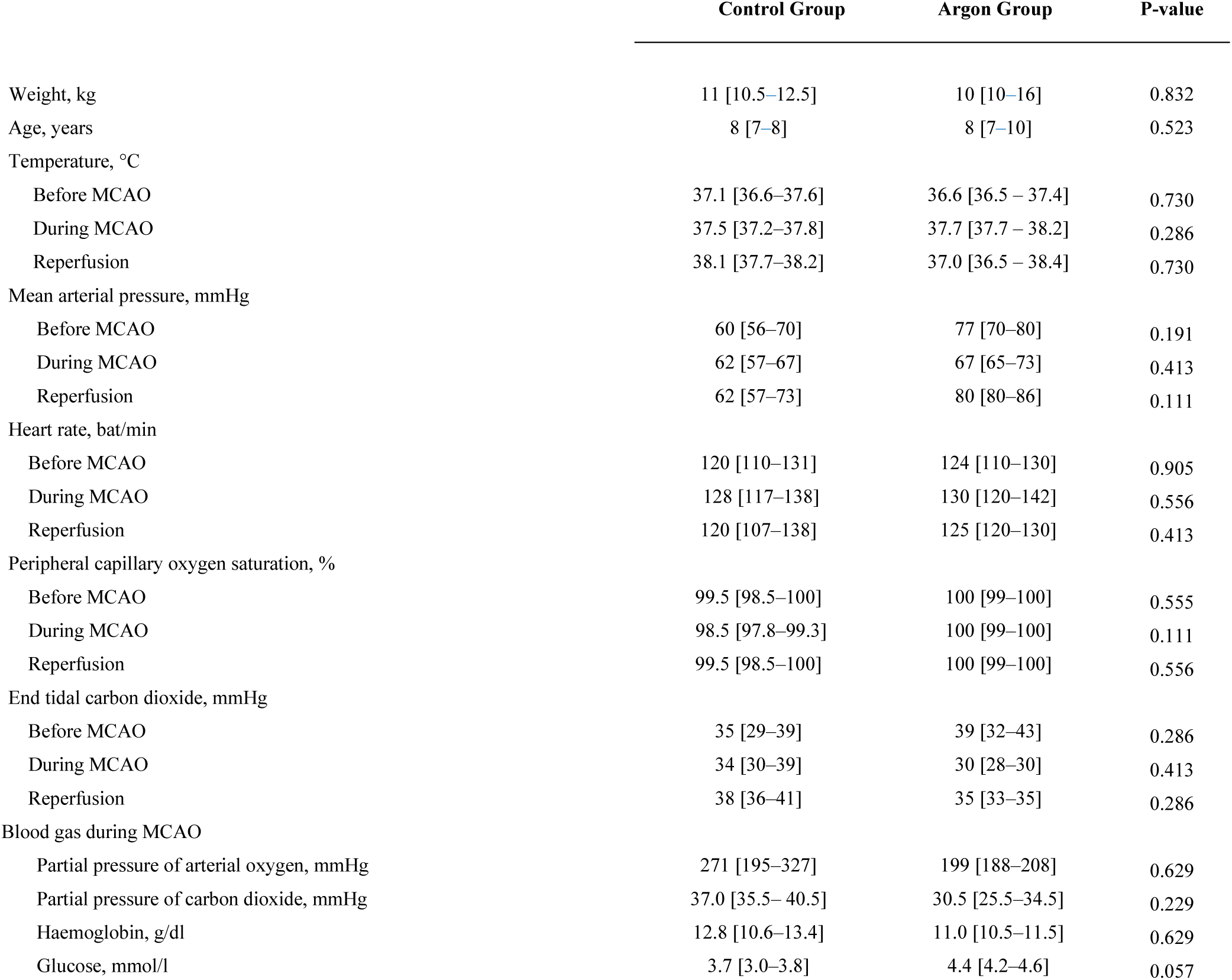
Physiologic Variables in Experimental Groups.

The density maps in Figure 4 allow clearer visualization of the distribution of brain damage. In the control group, the territories mostly affected by the ischemic core and edema were the supramarginal gyrus, proximal part of the superior temporal gyrus, postcentral gyrus, angular gyrus, and annectant gyrus (Figure 4A, C). In a small proportion of animals, the damage extended subcortically to the basal ganglia. The affected territories were more localized in the argon group (Figure 4B, D) and dominant in the proximal part of the superior temporal gyrus and underlying subcortical structures.

Eleven monkeys were used for the neurological clinical evaluation (Figure 5). One animal from the control group prematurely died because of a failure in the femoral artery catheterization leading to hemorrhagic shock. Therefore, it was not included in this analysis. Overall, the neurological assessments suggest better neurological prognosis in argon-treated animals than in control animals.

## DISCUSSION

The present study investigated the potential neuroprotective effect of the noble gas argon in a model of transient cerebral ischemia for the first time in NHPs. To this end, we used MRI scans acquired after brain reperfusion to compare ischemic core and edema volumes between a control group and a group of animals exposed to argon during the tMCAO procedure.

Based on the work of Zhao et al.[16] and with the objective of limiting the variability in ischemic core volumes and their location, we chose an endovascular approach for reversible occlusion of the middle cerebral artery. This approach is particularly suited to the objectives of our study for two main reasons. First, unlike surgical approaches for the placement of clips or ligatures, the endovascular approach is minimally invasive and preserves the integrity of the cranial cavity. Thus, it allows the pathophysiological consequences of cerebral ischemia, to be recreated as accurately as possible. Second, by preserving the cranium, this approach is particularly compatible with the assessment of brain damage by MRI and facilitates post-stroke neurological assessment. In the present study, we were able to perform a successful tMCAO procedure on 13 rhesus macaques with complete restoration of cerebral blood flow following ischemia. In keeping with middle cerebral artery occlusion in humans, this procedure led to ischemic core territories in the parieto-temporal cortical areas in our control group, which extended to the subcortical structures of the basal ganglia and internal capsule in some subjects[17]. As reported by others, the ischemic core volumes showed significant variability, which is consistent with the inter-subject variability of vascular collateralization in old world primates and humans[18].

Furthermore, NHP, particularly old-world monkeys, are better models for stroke research than rodents because they have more similar brain structure and function to humans[18,19]. The brains of NHP are more similar in size and complexity to the human brain, and they have a more developed cerebral cortex in proportion to subcortical structures[20]. In addition to having a more similar brain structure to humans compared to rodents, NHP also have more similar vascular brain organization and blood-brain barrier function[21]. These factors can be important in the context of stroke research, as they have a direct effect on the distribution and effectiveness of therapeutic agents in the brain. Furthermore, NHP have a more complex behavioral repertoire, which allows for a more detailed assessment of neurological outcome after stroke and is critical for understanding the long-term effects of a stroke on brain function and recovery[22].

Using our NHP model of tMCAO, we were able to demonstrate a significant effect of argon neuroprotection on the ischemic core volume. Moreover, we observed a better neurological outcome in neuroprotected animals, but this observation remains qualitative due to the limited size of our experimental groups. Overall, our results confirm the neuroprotective effects of argon inhalation observed in rat models of MCAO in terms of both infarct volumes[11,12,14] and neurological dysfunction[23,24]. However, in contrast to these rodent studies, we showed that the neuroprotective effect is limited to the cortical sheet, which was mainly affected in our NHP model of tMCAO. This difference is most likely related to structural differences between the rat and primate brain, particularly in the cortical and subcortical distribution of collateral arteries. The findings allow us to conclude positively on the neuroprotective effect of argon in cerebral ischemia, but further studies are needed to show that argon also has neuroprotective effects on brain damage following cerebral hemorrhage, cardiac arrest, or traumatic brain injuries as previously demonstrated in rodents[25–27].

Our study also confirms that, in terms of neuroprotection, argon is an alternative to xenon with two key advantages for the potential management of acute stroke patients. Firstly, thrombolysis currently remains a main method of restoring cerebral blood flow following stroke. Xenon, on the contrary, has been found to have an inhibitory effect on thrombolysis treatments[28]. Argon has no inhibitory effect on rtPA, and the in vitro potentiation of rtPA activity is controversial[29]. In line with these observations, we were able to successfully use a thrombolytic agent to restore cerebral blood flow in our animals in which a thrombus remained after the microcoil removal. Secondly, at atmospheric pressure, argon is not considered to have significant sedative properties, whereas xenon has been used as a general anesthetic and is known to have potent anesthetic effects through its action on the N-methyl-D-aspartate (NMDA) receptors[28]. From an experimental point of view, the lack of sedative properties of argon is critical because animal models of tMCAO, such as our NHP model, are performed under general anesthesia. In terms of the translational perspective, the lack of sedative effect of argon is particularly critical for the management of stroke patients with clinical approaches, such as thrombectomy, which require the use of general anesthesia or sedation in most of cases.

We specifically chose adult male monkeys because experimental studies show that young females may be protected against ischemic strokes by hormone-dependent mechanisms[30].

Notably, there are also limitations to using NHP in stroke research. NHP studies are more expensive and time-consuming to conduct than rodent studies, and they raise specific ethical considerations. These limitations must be considered when interpreting our results.

The first limitation concerns the number of animals used in this type of project. Based on Zhao et al.[16], who used a comparable model of tMCAO, we anticipated a median infarct volume of 2.2±0.7 cm^3^ in our control group and a 40% reduction in the infarcted brain volume relative to rodent studies. The experimentally observed volumes were larger and more variable (8.5±8.1 cm^3^) but, due to a strong effect of neuroprotection, these volumes were significantly different between our two experimental groups. However, the limited size of our groups combined with the loss of three animals in the control group did not allow us to make firm conclusions on the effect of neuroprotection on the neurological outcome of the tMCAO procedure. The second limitation is also related to the total number of animals involved in the project, which impacts the possibility of evaluating the neuroprotective effect of argon at different concentrations and at different timings with respect to ischemia onset. Rodent studies suggest that the neuroprotective effect of argon is dose-dependent, with a maximum efficacy at concentrations ranging from 50%[11] to 70%[26]. For our study, we favored a 60% argon concentration, which is in the middle these optimal values, and a favorable concentration to translate into a translational clinal set-up, allowing a balancing concentration of O2 of 40%, which cover the needs of most patients during mechanical thrombectomy. Regarding the timing of inhalation, we sought to recreate the conditions for rapid management of an ischemic stroke as realistically as possible, which, in the best-case scenario, could be initiated 30 minutes after ischemia onset. We also assumed that this time window would be optimal with respect to our own observations in a rat model of tMCAO in which pre-conditioning with argon initiated during ischemia provided an earlier and better level of neuroprotection than conditioning initiated before or after ischemia[23].

Finally, the interpretation of our results could be biased by the fact that the tMCAO procedure was performed under deep sevoflurane anesthesia, as this anesthetic has been suggested to have neuroprotective properties[31]. There are two counterarguments to this limitation. First, we were able to produce consistent ischemic cores in the animals of our control group even though they were under deep anesthesia before, during, and up to 3 hours after the tMCAO procedure. Second, several clinical studies have shown that maintaining deep anesthesia for thrombectomy compared to light sedation does not significantly improve the long-term neurological outcome of ischemic stroke patients[32,33].

In conclusion, this work provides the first evidence of the neuroprotective effect of argon inhalation in an *in vivo* model of tMCAO in NHPs. Our MRI investigations show a consistent neuroprotective effect of this noble gas in the acute phase of stroke with no adverse effects. Combined with the results obtained in *in vivo* investigations of argon neuroprotection in rodent models of tMCAO, this study makes the implementation of clinical tests in humans conceivable. This clinical phase will provide complementary information on the long-term persistence of argon neuroprotection and its consequences on functional recovery.

## Disclosures

The authors report no conflicts of interest.

**Supplementary figure 1. MRI analysis pipeline**: In the first processing step, if multiple T1w and T2w images were available, they were aligned and averaged for each modality. The T2w average was then co-registered to the T1w average. Both files were cropped and aligned, and a binary mask generated. The resulting images were denoised. The intensity inhomogeneity of T1 and T2 was corrected by applying an N4 debias correction, and bias correction was computed on T1w using the product of the T1w and T2w intensities. The masked T1w images of each subject were then normalized to the INIA19 template[34]. The second step of the pipeline segmented the T1 from the tissues. Images were segmented into four types: cerebrospinal fluid (CSF), white matter (WM), grey matter (GM), and dark non-brain matter using the SPM12 old segment method. White matter tissue projected back in native space was then used for normalization of b0 mean to T2 (see diffusion part). The third step processed diffusion-tensor imaging (DTI). Top-up estimated the susceptibility-induced off-resonance field from pairs of images, with reversed phase-encode blips and distortions going in opposite directions. Two non–diffusion-weighted images with opposed phase-encoding polarities were extracted, and susceptibility-induced distortion was estimated top-up. The eddy tool corrects image distortions by assuming that diffusion signals obtained from two MPG directions with a small angle difference are similar, combining the correction for susceptibility and eddy currents/movements. These diffusion images were processed with the FSL tool dtifit[35]. B0 images were averaged and aligned on the T2, and then normalized to the INIA19 template space, resulting in the aDC image normalized to the INIA19 space, and the normalized aDC was used to mark the ischemic core for each subject. Finally, two masks were applied to separately quantify marked voxels from the cortical or subcortical areas (Supplementary Figure 2).

**Supplementary figure 2. Cortical and subcortical masks:** (A) 3D volume, (B) axial, (C) sagittal, and (D) coronal views of the masks that were used to separately quantify cortical (yellow) and subcortical (red) damage. The cortical mask includes the following territories: occipital gyrus, occipital white matter, lingual gyrus, cuneus, inferior, occipital gyrus, annectant gyrus, fusiform gyrus, angular gyrus, precuneus, superior parietal lobule, inferior temporal gyrus, supramarginal gyrus, inferior olivary complex, occipital horn of the lateral ventricle, superior temporal gyrus, cerebral white matter, middle temporal gyrus, fasciolar gyrus, posterior cingulate gyrus, posterior parahippocampal gyrus, isthmus of the cingulate gyrus, splenium of the corpus callosum, dentate gyrus, postcentral gyrus, body of the corpus callosum, precentral gyrus, claustrum, extreme capsule, temporal white matter, insula, entorhinal area, anterior cingulate gyrus, frontal white matter, superior frontal gyrus, olfactory tract, middle frontal gyrus, inferior frontal gyrus, fronto-orbital gyrus, lateral orbital gyrus, straight gyrus, medial orbital gyrus, central sulcus, area 8 alpha of Vogts, area 23 of Mauss, area PrCO of Roberts, cingulum limitans, parasubicular area, prosubiculum, subiculum, presubiculum, endopiriform nucleus, arachnoid mater, callosal sulcus, olfactory bulb, and pars anterior of the paramedian lobule. All the remaining territories were included in the subcortical mask.

## REFERENCES

1. Lopez AD, Mathers CD, Ezzati M, Jamison DT, Murray CJ: Global and regional burden of disease and risk factors, 2001: systematic analysis of population health data. The Lancet 2006; 367:1747–57

2. Roger VL, Go AS, Lloyd-Jones DM, Adams RJ, Berry JD, Brown TM, Carnethon MR, Dai S, Simone G de, Ford ES, Fox CS, Fullerton HJ, Gillespie C, Greenlund KJ, Hailpern SM, Heit JA, Ho PM, Howard VJ, Kissela BM, Kittner SJ, Lackland DT, Lichtman JH, Lisabeth LD, Makuc DM, Marcus GM, Marelli A, Matchar DB, McDermott MM, Meigs JB, Moy CS, et al.: Heart Disease and Stroke Statistics—2011 Update: A Report From the American Heart Association. Circulation 2011; 123

3. Paul S, Candelario-Jalil E: Emerging neuroprotective strategies for the treatment of ischemic stroke: An overview of clinical and preclinical studies. Exp Neurol 2021; 335:113518

4. Hernández-Jiménez M, Abad-Santos F, Cotgreave I, Gallego J, Jilma B, Flores A, Jovin TG, Vivancos J, Molina CA, Montaner J, Casariego J, Dalsgaard M, Hernández-Pérez M, Liebeskind DS, Cobo E, Ribo M: APRIL: A double-blind, placebo-controlled, randomized, Phase Ib/IIa clinical study of ApTOLL for the treatment of acute ischemic stroke. Front Neurol 2023; 14:1127585

5. Rizvi M, Jawad N, Li Y, Vizcaychipi MP, Maze M, Ma D: Effect of noble gases on oxygen and glucose deprived injury in human tubular kidney cells. Exp Biol Med 2010; 235:886–91

6. Zhuang L, Yang T, Zhao H, Fidalgo AR, Vizcaychipi MP, Sanders RD, Yu B, Takata M, Johnson MR, Ma D: The protective profile of argon, helium, and xenon in a model of neonatal asphyxia in rats*: Crit Care Med 2012; 40:1724–30

7. Cattano D, Williamson P, Fukui K, Avidan M, Evers AS, Olney JW, Young C: Potential of xenon to induce or to protect against neuroapoptosis in the developing mouse brain. Can J Anesth Can Anesth 2008; 55:429–36

8. Liang M, Ahmad F, Dickinson R: Neuroprotection by the noble gases argon and xenon as treatments for acquired brain injury: a preclinical systematic review and meta-analysis. Br J Anaesth 2022; 129:200–18

9. Růžička J, Beneš J, Bolek L, Markvartová V: Biological effects of noble gases. Physiol Res 2007:S39–44 doi:10.33549/physiolres.931300

10. Ulbrich F, Goebel U: The Molecular Pathway of Argon-Mediated Neuroprotection. Int J Mol Sci 2016; 17:1816

11. David HN, Haelewyn B, Degoulet M, Colomb DG, Risso J-J, Abraini JH: Ex Vivo and In Vivo Neuroprotection Induced by Argon When Given after an Excitotoxic or Ischemic Insult. PLoS ONE Edited by Arai K. 2012; 7:e30934

12. Ryang Y-M, Fahlenkamp AV, Rossaint R, Wesp D, Loetscher PD, Beyer C, Coburn M: Neuroprotective effects of argon in an in vivo model of transient middle cerebral artery occlusion in rats*: Crit Care Med 2011; 39:1448–53

13. Gardner AJ, Menon DK: Moving to human trials for argon neuroprotection in neurological injury: a narrative review. Br J Anaesth 2018; 120:453–68

14. Fahlenkamp AV, Coburn M, Prada A de, Gereitzig N, Beyer C, Haase H, Rossaint R, Gempt J, Ryang Y-M: Expression analysis following argon treatment in an in vivo model of transient middle cerebral artery occlusion in rats. Med Gas Res 2014; 4:11

15. Wu D, Wu L, Chen J, Huber M, He X, Li S, Ding Y, Ji X: Primate Version of Modified Rankin Scale for Classifying Dysfunction in Rhesus Monkeys. Stroke 2020; 51:1620–3

16. Zhao B, Shang G, Chen J, Geng X, Ye X, Xu G, Wang J, Zheng J, Li H, Akbary F, Li S, Lu J, Ling F, Ji X: A more consistent intraluminal rhesus monkey model of ischemic stroke. Neural Regen Res 2014; 9:2087

17. Phan TG, Donnan GA, Wright PM, Reutens DC: A Digital Map of Middle Cerebral Artery Infarcts Associated With Middle Cerebral Artery Trunk and Branch Occlusion. Stroke 2005; 36:986–91

18. Cook DJ, Teves L, Tymianski M: Treatment of stroke with a PSD-95 inhibitor in the gyrencephalic primate brain. Nature 2012; 483:213–7

19. Kwiecien TD, Sy C, Ding Y: Rodent models of ischemic stroke lack translational relevance… are baboon models the answer? Neurol Res 2014; 36:417–22

20. Passingham R: How good is the macaque monkey model of the human brain? Curr Opin Neurobiol 2009; 19:6–11

21. Lin X, Wang H, Chen J, Zhao P, Wen M, Bingwa LA, Jin K, Zhuge Q, Yang S: Nonhuman primate models of ischemic stroke and neurological evaluation after stroke. J Neurosci Methods 2022; 376:109611

22. Won J, Yi KS, Choi C-H, Jeon C-Y, Seo J, Kim K, Yeo H-G, Park J, Kim YG, Jin YB, Koo B-S, Lim KS, Lee S, Kim KJ, Choi WS, Park S-H, Kim Y-H, Huh J-W, Lee S-R, Cha S-H, Lee Y: Assessment of Hand Motor Function in a Non-human Primate Model of Ischemic Stroke. Exp Neurobiol 2020; 29:300–13

23. Giacomino L, Triglia T, Garrigue P, Brige P, Bruder N, Guillet B, Velly L: Effet neuroprotecteur à long terme de l’Argon sur un modèle in vivo d’ischémie-reperfusion cérébrale. Anesth Réanimation 2015; 1:A182–3

24. Liu J, Veldeman M, Höllig A, Nolte K, Liebenstund L, Willuweit A, Langen K-J, Rossaint R, Coburn M: Post-stroke treatment with argon preserved neurons and attenuated microglia/macrophage activation long-termly in a rat model of transient middle cerebral artery occlusion (tMCAO). Sci Rep 2022; 12:691

25. Höllig A, Schug A, Fahlenkamp A, Rossaint R, Coburn M, Argon Organo-Protective Network (AON): Argon: Systematic Review on Neuro- and Organoprotective Properties of an “Inert” Gas. Int J Mol Sci 2014; 15:18175–96

26. Brücken A, Kurnaz P, Bleilevens C, Derwall M, Weis J, Nolte K, Rossaint R, Fries M: Dose dependent neuroprotection of the noble gas argon after cardiac arrest in rats is not mediated by KATP—Channel opening. Resuscitation 2014; 85:826–32

27. Schneider FI, Krieg SM, Lindauer U, Stoffel M, Ryang Y-M: Neuroprotective Effects of the Inert Gas Argon on Experimental Traumatic Brain Injury In Vivo with the Controlled Cortical Impact Model in Mice. Biology 2022; 11:158

28. David HN, Haelewyn B, Risso J-J, Colloc’h N, Abraini JH: Xenon is an Inhibitor of Tissue-Plasminogen Activator: Adverse and Beneficial Effects in a Rat Model of Thromboembolic Stroke. J Cereb Blood Flow Metab 2010; 30:718–28

29. David HN, Haelewyn B, Risso J-J, Abraini JH: Modulation by the noble gas argon of the catalytic and thrombolytic efficiency of tissue plasminogen activator. Naunyn Schmiedebergs Arch Pharmacol 2013; 386:91–5

30. Jiang M, Ma C, Li H, Shen H, Li X, Sun Q, Chen G: Sex Dimorphisms in Ischemic Stroke: From Experimental Studies to Clinic. Front Neurol 2020; 11:504

31. Canas PT, Velly LJ, Labrande CN, Guillet BA, Sautou-Miranda V, Masmejean FM, Nieoullon AL, Gouin FM, Bruder NJ, Pisano PS: Sevoflurane Protects Rat Mixed Cerebrocortical Neuronal–Glial Cell Cultures against Transient Oxygen–Glucose Deprivation. Anesthesiology 2006; 105:990–8

32. Schönenberger S, Uhlmann L, Hacke W, Schieber S, Mundiyanapurath S, Purrucker JC, Nagel S, Klose C, Pfaff J, Bendszus M, Ringleb PA, Kieser M, Möhlenbruch MA, Bösel J: Effect of Conscious Sedation vs General Anesthesia on Early Neurological Improvement Among Patients With Ischemic Stroke Undergoing Endovascular Thrombectomy: A Randomized Clinical Trial. JAMA 2016; 316:1986

33. Maurice A, Eugène F, Ronzière T, Devys J-M, Taylor G, Subileau A, Huet O, Gherbi H, Laffon M, Esvan M, Laviolle B, Beloeil H, for the GASS (General Anesthesia versus Sedation for Acute Stroke Treatment) Study Group and the French Society of Anesthesiologists (SFAR) Research Network: General Anesthesia *versus* Sedation, Both with Hemodynamic Control, during Intraarterial Treatment for Stroke: The GASS Randomized Trial. Anesthesiology 2022; 136:567–76

34. Rohlfing T, Kroenke CD, Sullivan EV, Dubach MF, Bowden DM, Grant KA, Pfefferbaum A: The INIA19 Template and NeuroMaps Atlas for Primate Brain Image Parcellation and Spatial Normalization. Front Neuroinformatics 2012; 6

35. Yamada H, Abe O, Shizukuishi T, Kikuta J, Shinozaki T, Dezawa K, Nagano A, Matsuda M, Haradome H, Imamura Y: Efficacy of Distortion Correction on Diffusion Imaging: Comparison of FSL Eddy and Eddy_Correct Using 30 and 60 Directions Diffusion Encoding. PLoS ONE Edited by Najbauer J. 2014; 9:e112411

