## Supplementary figures and images for "Argon neuroprotection in a non-human primate model of transient endovascular ischemic stroke"

### Supplemental Figure 1

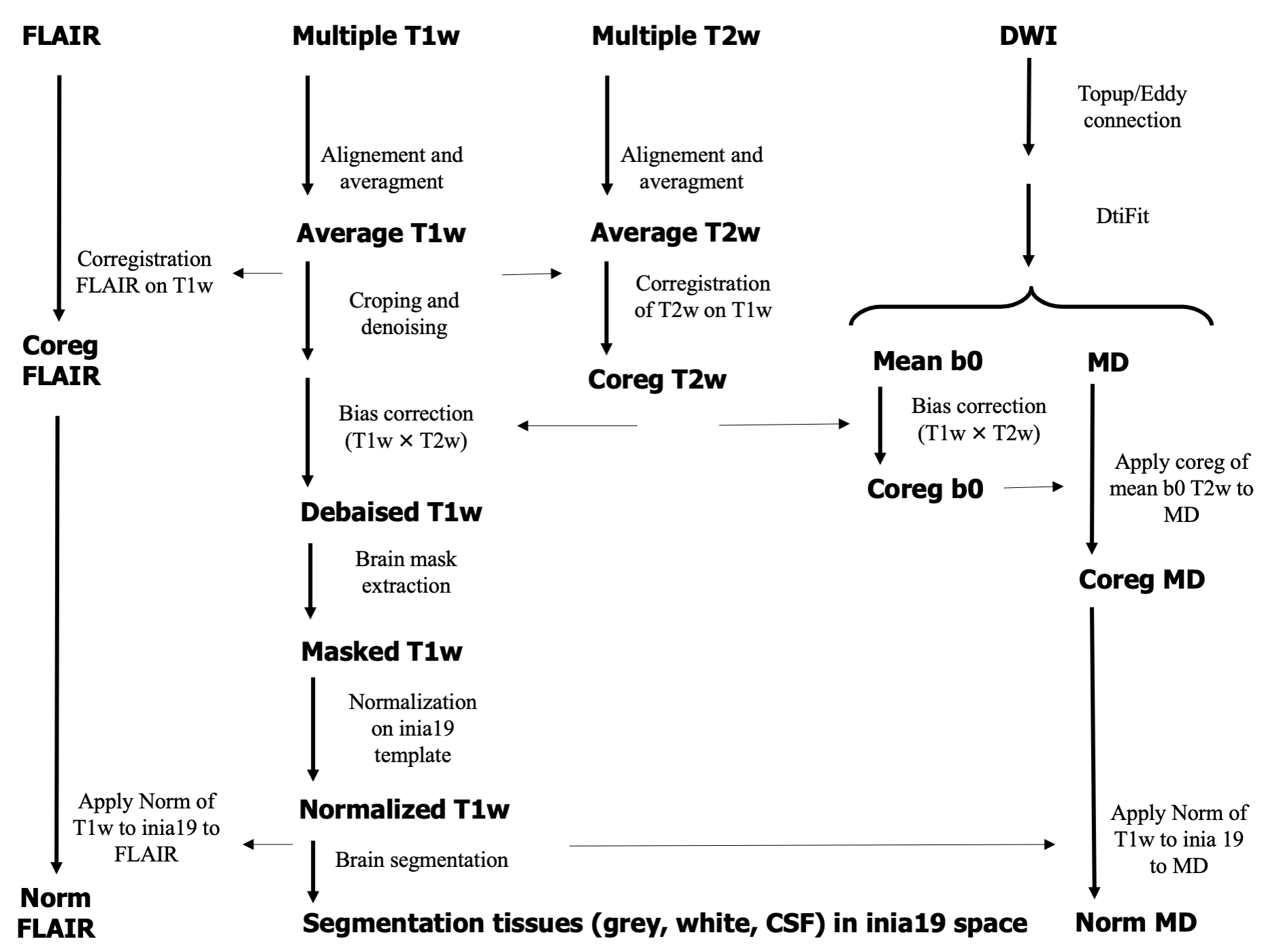

### Supplemental Figure 2

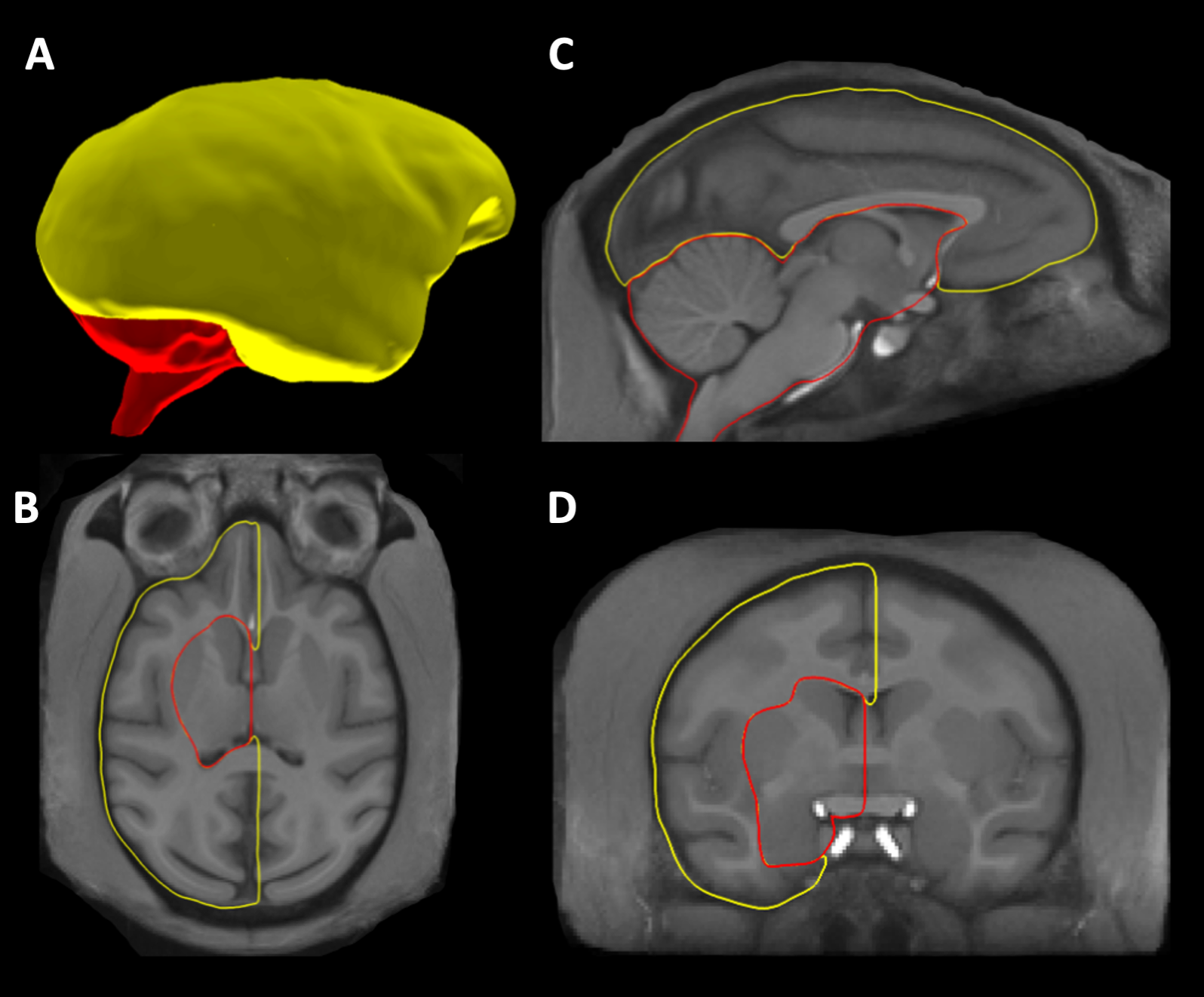
